# A pigtailed macaque model for Kyasanur Forest disease virus and Alkhurma hemorrhagic disease virus

**DOI:** 10.1101/2021.06.01.446549

**Authors:** Rebecca M. Broeckel, Friederike Feldmann, Kristin L. McNally, Abhilash I. Chiramel, Patrick W. Hanley, Jamie Lovaglio, Rebecca Rosenke, Dana P. Scott, Greg Saturday, Fadila Bouamr, Shelly J. Robertson, Sonja M. Best

## Abstract

Kyasanur Forest disease virus (KFDV) and the closely related Alkhurma hemorrhagic disease virus (AHFV) are emerging flaviviruses that cause severe viral hemorrhagic fevers in humans. Increasing geographical expansion and case numbers, particularly of KFDV in southwest India, class these viruses as a public health threat. Viral pathogenesis is not well understood and additional vaccines and antivirals are needed to effectively counter the impact of these viruses. However, current animal models for KFDV do not accurately reproduce viral tissue tropism or clinical outcomes observed in humans. Here, we show pigtailed macaques (*Macaca nemestrina*) infected with KFDV or AHFV develop viremia that peaks 2 to 4 days following inoculation. Over the course of infection, animals developed lymphocytopenia, thrombocytopenia, and elevated liver enzymes. Infected animals exhibited hallmark signs of human disease characterized by a flushed appearance, piloerection, dehydration, loss of appetite, weakness, and hemorrhagic signs such as epistaxis. Virus was commonly present in the gastrointestinal tract, consistent with human disease caused by KFDV and AHFV where gastrointestinal symptoms (hemorrhage, vomiting, diarrhea) are common. This work characterizes a nonhuman primate model for KFDV and AHFV that closely resembles human disease for further utilization in understanding host immunity and development of antiviral countermeasures.

**Author Summary:** Kyasanur Forest disease virus (KFDV) and Alkhurma hemorrhagic disease virus (AHFV) are tick-borne flaviviruses that cause viral hemorrhagic fevers in India and the Arabian Peninsula, respectively. Bonnet macaques and black-faced langurs are susceptible to KFDV infection, but these animals do not experience hemorrhagic signs as seen in human cases with KFDV. This work characterizes for the first time experimental infection of KFDV and AHFV in pigtailed macaques (PTMs). Infected PTMs can develop moderate to severe disease that mirrors many aspects of human disease, including some hemorrhagic signs. Together these data describe the PTM model for KFDV and AHFV as a valuable tool for future work to study viral pathogenesis and for assessing the efficacy of vaccines and antivirals.

## Introduction

Kyasanur Forest disease virus (KFDV) and Alkhurma hemorrhagic fever virus (AHFV) are emerging flaviviruses that can cause fatal hemorrhagic disease in humans. These viruses are NIAID category C priority pathogens that require a maximum containment facility to conduct research. Transmission of these viruses to humans primarily occurs following the bite of infected ticks. KFDV was originally discovered in the Shimoga district of Karnataka but has expanded its geographical distribution in the last decade to include Kerala, Goa, and Maharashtra states (1). An estimated 100-900 cases of KFDV occur annually with a 2-10% case-fatality rate (1-3). KFDV causes a sudden onset of acute febrile illness including headache, fever, vomiting, diarrhea, inflammation of the eyes, and dehydration. Disease may progress to include hemorrhagic signs particularly bleeding from nose, gums, gastrointestinal tract, or lungs (3). AHFV is a newly emerging genetic variant of KFDV that was first isolated in 1995 (4, 5). While KFDV is localized to southwest India, AHFV cases were identified in Egypt and Saudi Arabia (4, 6). AHFV causes similar febrile illness and hemorrhagic disease in humans as observed with KFDV (7). However, very little is known regarding mechanisms of disease.

The expanding geographic range of tick-borne flaviviruses (TBFVs) witnessed over the last several decades, the steady increase in annual cases, and the potential for severe clinical disease necessitates the development of relevant animal models to study viral pathogenesis and to evaluate vaccines and antiviral therapies (1, 8, 9). Small animal models for TBFVs have limitations in that they do not recapitulate key aspects of human disease. Several commonly available strains of laboratory mice are susceptible to KFDV and AHFV infection, with mice developing neurological signs 5-10 days after infection associated with high virus burden in the brain at clinical endpoints (10-14). However, the neurotropic nature of KFDV and AHFV in the mouse model does not recapitulate the hemorrhagic nature of human disease. Other commonly used small animal models (ferrets, guinea pigs, and hamsters) do not show signs of disease following infection with KFDV, despite the presence of virus in the brain of infected hamsters (15).

Non-human primates (NHPs) have been investigated as potential models of TBFV pathogenesis aith susceptibility to infection being highly species-dependent. Early reports suggested that rhesus macaques (*Macaca mulatta)* infected with KFDV may develop viremia but clinical signs of disease were absent (3, 16). Red-faced bonnet macaques (*Macaca radiata*) and black-faced langurs (*Presbytis entellus*) are naturally susceptible to KFDV, and clusters of KFDV human cases coincide with discovery of deceased bonnet macaques and langurs in local forested areas (3). Experimental infection of these two NHP species demonstrated that black-faced langurs in particular are highly susceptible to KFDV and succumbed to infection during the viremic phase (17). Bonnet macaques infected with KFDV experience a biphasic illness with 10-100% lethality depending on the study (17-20). The acute phase is characterized by dehydration, hypotension, bradycardia and diarrhea, while neurological signs such as tremors may be observed in the second phase (17, 18, 20, 21). However, platelets were only mildly decreased over infection, and no hemorrhagic signs were reported in these animals (18, 19). Histopathological abnormalities in KFDV-infected bonnet macaques are typically observed in the gastrointestinal tract, lymph node, spleen, kidney, and liver (3, 18, 19). Compared to human KFDV disease, infected bonnet macaques develop the second neurological disease phase more frequently, with KFDV routinely isolated from brain and CSF samples (18, 21). In addition, while bonnet macaques and black-faced langurs have application to studying KFDV disease, these species are not readily available for biomedical research outside of India, thus preventing their wider use as a comparative model for KFDV.

In an effort to develop improved disease models, we investigated whether pigtailed macaques (PTMs) were susceptible to infection with KFDV and AHFV. PTMs have been used in development of pathogenesis models for HIV-1, Zika virus (a related flavivirus), influenza virus, Norwalk-like virus, and human herpesvirus 6A coinfections with SIV (22-27). KFDV- and AHFV-infected PTMs displayed moderate to severe clinical illness and some hemorrhagic signs. The PTM KFDV and AHFV models closely follow reports of KFDV and AHFV infection in humans and captures aspects of human disease that have not been described in other KFDV NHP models. Thus PTMs represent an important model for countermeasure development against these emerging viruses.

## Results

### Pilot studies of KFDV infection of PTMs

To determine whether PTMs are susceptible to infection with KFDV, a pilot experiment was performed where two animals were infected subcutaneously (sc) to mimic a tick bite between the shoulder blades with 10^5^ pfu KFDV P9605. Both KFDV-infected animals (KFDV 1 and KFDV 2) showed clinical signs of infection (Table S1), although these were not consistent between the two KFDV-infected animals (Figure S1). KFDV 1 had decreased appetite, piloerection, and hunched posture whereas KFDV 2 had decreased appetite and transient bloody nasal discharge. Animals had fully recovered by 14 dpi, and they did not develop any additional clinical signs through day 43. Infectious virus was not detected in the plasma of KFDV 1, but virus was directly isolated from the plasma of KFDV 2 at the peak of viremia at 4 dpi. Neutralizing antibodies were detected in KFDV-infected animals starting at 8-10 dpi. TBFV-specific antibodies are known to cross-neutralize other TBFVs in culture and may protect against disease caused by heterologous TBFVs *in vivo* (11, 28). Therefore, we tested whether 28 dpi immune sera from KFDV sc-inoculated animals could neutralize AHFV and TBEV by a plaque reduction neutralization assay. A dilution of 1:100 of sera could neutralize over 50% of the AHFV and TBEV plaques (Figure S2), which is consistent with previous reports of cross-neutralizing activity of TBFV immune sera. No significant lesions were noted at necropsy, except KFDV 1 had a pale, reticulated liver (Table 1). This pilot study suggested that PTMs are susceptible to KFDV infection and have the potential to develop viremia and disseminated infection.

**Table 1.** KFDV and AHFV Necropsy findings.

|  | KFDV 1 (sc) | KFDV 2 (sc) | KFDV 3 (sc/iv) | KFDV 4 (sc/iv) | KFDV 5 (sc/iv) | KFDV 6 (sc/iv) | KFDV 7 (sc/iv) | KFDV 8 (sc/iv) | AHFV 1 (sc/iv) | AHFV 2 (sc/iv) | AHFV 3 (sc/iv) | AHFV 4 (sc/iv) |
| --- | --- | --- | --- | --- | --- | --- | --- | --- | --- | --- | --- | --- |
| Day post infection | D42 | D42 | D6 | D8 | D8 | D9 | D9 | D7 | D9 | D9 | D9 | D8 |
| Discharge | Normal | Normal | Normal | Epistaxis | Epistaxis | Dried blood in nasal cavity | None | Epistaxis | None | Dried blood in nasal cavity | Dried blood in nasal cavity | None |
| Subcutis | Normal | Normal | Normal | Tacky | Normal | Tacky | Normal | Tacky | Tacky | Normal | Tacky | Edema, Tacky |
| Liver | Pale | Normal | Yellow-orange | Pale | Pale | Pale | Pale tan | Pale | Pale | Normal | Normal | Pale |
| Liver Spleen | Reticulated |  | Fatty | Friable | Reticulated | Enlarged | Enlarged | Reticulated | Enlarged |  |  | Enlarged |
|  |  |  | Reticulated | Fatty | Fatty | Rounded Edges | Swollen | Prominent Vasculature | Rounded Edges |  |  | Lumpy |
|  |  |  |  |  |  |  |  |  | Multifocal Hemorrhages |  |  |  |
|  | Normal | Normal | Adhesions to mesenteric | Enlarged, firm | Normal | Normal | Normal | Normal | Lumpy Irregular surface | Normal | Enlarged | Enlarged, Turgid |
| Lungs | Normal | Normal | All lobes hyperemic | Congested | Normal | Multifocal <1mm black spots | Adhesions | Normal | Hemorrhagic, dorsal ecchymosis all lobes, caudal lobes ventral surface | Failure to collapse | Normal | Small bulla on R lower lung lobe |
| Lungs Adhesions |  |  |  | Left lung all lobes hyperemic |  |  |  |  |  |  |  |  |
|  | Normal | Normal | Adhesions throughout abdomen | Mesenteric adhesions | Normal | Normal | Adhesions in chest and lung cavity | Multiple Adhesions lower abdomen | Normal | Normal | Normal | Normal |
| Other Comments |  |  |  | Jejunum fluid-filled | Large parotid salivary glands |  |  | Mediastinal LN edematous |  |  | Dehydrated | Dehydrated |
|  |  |  |  | Foam in trachea |  |  |  | Gall bladder distended |  |  |  |  |

### Clinical signs and tissue distribution of virus in KFDV-infected PTMs following sc/iv inoculation

Due to the variable clinical signs and viremia between individual animals in the pilot study, an additional study was conducted using a combination of subcutaneous and intravenous (sc/iv) inoculation with KFDV P9605. In total, six PTMs (KFDV 3-8) were infected and monitored for signs of illness (Figure 1). Clinical signs included severely decreased appetite, piloerection, flushed appearance, dehydration, and cyanotic mucous membranes (Table 2). Epistaxis was observed in 5 out of 6 animals, starting at 6 dpi and continuing through the end of the experiment. Two animals (KFDV 3 and KFDV 4) had irregular breathing patterns observable from 4 dpi. Four of the six animals met the pre-determined euthanasia criteria of a clinical score of 35 or greater between 6-8 dpi. The clinical scoring of the remaining two animals did not reach a score of 35 and was beginning to resolve at 9 dpi, prompting euthanasia for virus isolation from tissues. All six animals had demonstrable viremia with infectious virus isolated between 2-6 dpi, with peak titer at day 4 (Figure 1C). Neutralizing antibodies were detectable in serum at 8 dpi at a dilution of 1:100 (Figure 1D). Neutralizing activity was not detected in terminal sera from KFDV 3 and KFDV 8, that reached euthanasia criteria at 6 and 7 dpi, respectively (data not shown). All oral swabs taken on exam days were negative for KFDV. However, infectious virus was isolated from rectal swabs in two out of six animals on days 4 and/or 6 post infection suggesting virus shedding from the gastrointestinal tract.

**Figure 1.**
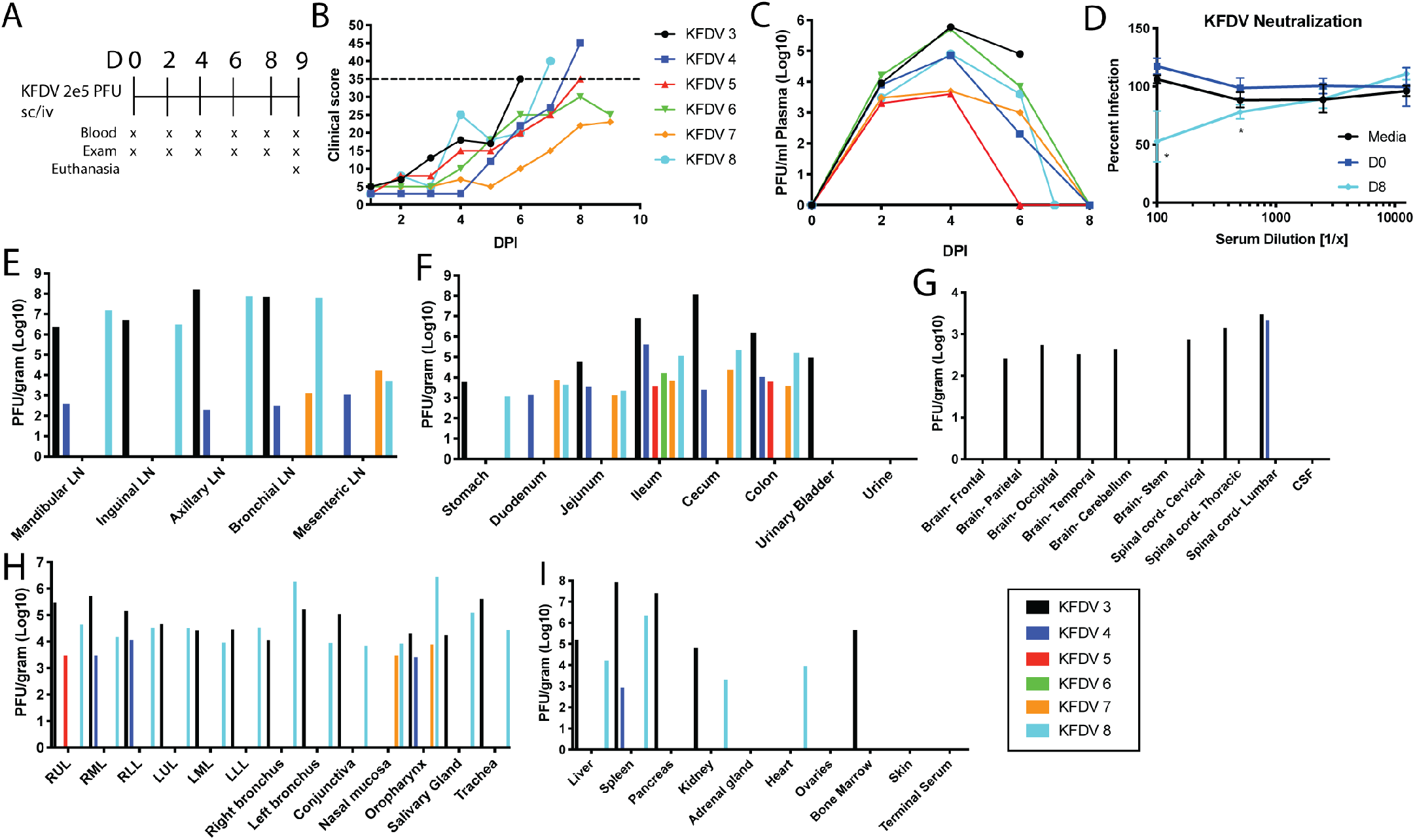
KFDV causes moderate to severe clinical illness in pigtailed macaques, and virus is present in multiple tissues. Six PTMs (KFDV 3-8) were infected with 2 × 10^5^ pfu KFDV P9605 sc/iv (10^5^ pfu, per route). **A**. Exam schedule following infection with KFDV. **B**. Animals were scored daily for clinical signs following infection. The dotted line indicates a score of 35. **C**. KFDV viremia was measured by limiting dilution plaque assay from plasma on the indicated days post infection. **D**. Neutralization assays were performed using heat-inactivated serum collected at 8 dpi by plaque assay. Media indicates no sera. Data are normalized to D0. Error bars represent the standard deviation (SD) across four animals. Statistics were performed using Sidak’s multiple comparison test, comparing D0 to D8 at all serum dilutions, (*N* = 4; \**P* < 0.05). **E-I**. At terminal endpoints, or 9 dpi, animals were euthanized and virus was isolated from various tissues by plaque assay. Plaque counts were normalized to 1 gram of tissue. Data from each individual animal are plotted.

**Table 2.** KFDV and AHFV clinical observations. Pigtailed macaques were observed for clinical signs following sc/iv inoculation of KFDV or AHFV. Some parameters (mucous membrane color, dehydration, breathing quality, CRT, and temperature) were only measured during exams. The presence or absence of a clinical sign are reported as Y (yes) or N (no).

|  | KFDV 3<br>(sc/iv) | KFDV 4<br>(sc/iv) | KFDV 5<br>(sc/iv) | KFDV 6<br>(sc/iv) | KFDV 7<br>(sc/iv) | KFDV 8<br>(sc/iv) | AHFV 1<br>(sc/iv) | AHFV 2<br>(sc/iv) | AHFV 3<br>(sc/iv) | AHFV 4<br>(sc/iv) | Frequency<br>(sc/iv) n =<br>10 |
| --- | --- | --- | --- | --- | --- | --- | --- | --- | --- | --- | --- |
| Decreased appetite | Severe | Severe | Severe | Severe | Decreased | Severe | Decreased | Decreased | Decreased | Severe | 100% |
| Loss of interest in treats | Y | N | Y | Y | N | Y | N | N | N | Y | 50% |
| Piloerection | Y | N | Y | Y | Y | Y | Y | N | N | Y | 70% |
| Hunched posture | N | Y | Y | Y | Y | N | Y | N | N | N | 50% |
| Loose stool | N | N | N | N | N | N | N | N | N | N | 0% |
| No feces | Y | N | N | N | N | N | N | N | N | N | 10% |
| Clear nasal discharge | N | N | N | Y | N | Y | N | N | N | N | 20% |
| Transient epistaxis | N | D8 heavy epistaxis | Y | Y | Y | Y | Dried blood on nose | Y | Y | N | 80% |
| Unsteady | N | Y | N | N | N | Y | N | N | N | N | 20% |
| Slow and careful movements | N | Y | Y | N | N | Y | Y | N | N | N | 40% |
| Reluctant to move | Y | N | Y | N | N | N | N | N | N | N | 20% |
| Using cage for support | Y | Y | N | N | N | Y | Y | N | N | N | 40% |
| Mucous membranes | bluish D6 | bluish D6-8 | bluish D6, D8 | N | a little blood on gums | D4-7 Muddy | N | N | N | D8 Pale pink muddy | 100% |
| Dehydration | D6 5-8% | N | 5-10% | D4-6 5-8%, D9 8-10% | D9 5-8% | D4-7 5-8% | D6 5-8%, D9 10-12% | D4-9 5-8% | D6-9 10-12% | D4 5-8%, D8 10% | 90% |
| CRT 2 seconds or more | D6, 3 sec. | D8, 2 sec | N | N | N | N | N | N | N | N | 20% |
| Temperature change > 1 °C | N | Temp drop 1.6 °C D8 | Temp drop 1.7 °C D8 | N | Temp drop 1.5 °C D9 | N | Temp drop 1.6 °C D9 | Temp drop 1.2 °C D9 | Temp drop 1.7 °C D8 | Temp drop 1.7 °C D8 | 70% |
| Flushed Appearance | D6 | N | D6-8 | D6 | N | D6-7 | D6-8 | N | D4-8 | D7-8 pale/green | 70% |
| Mild Facial edema | N | N | N | N | N | N | N | N | Y | N | 10% |
| Other Comments |  |  |  | Liver palpable D6-9 |  |  |  |  |  |  |  |
| Score 35 | D6 | D8 | D8 | N | N | D7 | D9 | N | N | N | 50% |

Infectious virus was readily detected in several tissues from necropsied animals by plaque assay (Figure 1 E-I). The most consistent finding was the presence of virus in the gastrointestinal tract from all KFDV-infected animals. Infectious virus was recovered in the lungs, lymph nodes, and oropharynx in four of six animals. Two animals (KFDV 3 and KFDV 8) had virus present in almost every tissue examined. Virus was typically not present in the CNS, except for two animals, KFDV 3 and KFDV 4 that had virus present in the brain and spinal cord (KFDV 3) or the lumbar spinal cord (KFDV 4). Thus, PTMs infected with KFDV develop clinical illness with highest infectious virus burden in the gastrointestinal tract and lymphoid tissues from days 6-9 post infection, consistent with viral shedding detected in rectal swabs. Importantly, virus tropism to the gastrointestinal tissues in the PTM model accurately reflects human clinical disease, where gastrointestinal symptoms are a hallmark sign of KFDV infection.

### Clinical signs and tissue distribution of virus in AHFV-infected PTMs

AHFV is a genetic variant of KFDV, and causes a similar hemorrhagic disease in humans although its pathogenesis has not been examined in NHP models. Therefore, we extended the model by infecting four PTMs with AHFV 200300001 by the sc/iv route, followed by monitoring for clinical signs with exams and blood draws every two days (Figure 2A). AHFV-infected animals developed disease signs that resembled infection with KFDV, including a flushed appearance, dehydration, and transient epistaxis (Table 2). One out of four animals reached a clinical score of 35 on day 9 post infection, while the other animals had a maximum clinical score of 28 to 33 (Figure 2B). As it was clear from clinical parameters that the disease was resolving in the remaining three animals, all PTMs were euthanized at 8 or 9 dpi to examine virus tissue distribution (Figure 2E-I). All four animals had a detectable viremia between day 2 and 4 post infection (Figure 2C) and developed neutralizing antibodies by day 8 post infection at a 1:100 dilution of serum (Figure 2D). However, all rectal swabs taken on exam days were negative for infectious virus. The oral swab of AHFV 4 was positive on day 4 post infection, but all other oral swabs were negative. Despite a lack of shedding, AHFV could be isolated from the gastrointestinal tract in all four animals, similar to KFDV (Figure 2F). Three out of four animals also had virus in the lymph nodes, oropharynx, and lungs. No virus was detectable in the brain, CSF, or spinal cord (Figure 2E-I). Thus, PTMs develop clinical signs of illness and virus can be isolated from multiple tissues following infection with either KFDV or AHFV.

**Figure 2.**
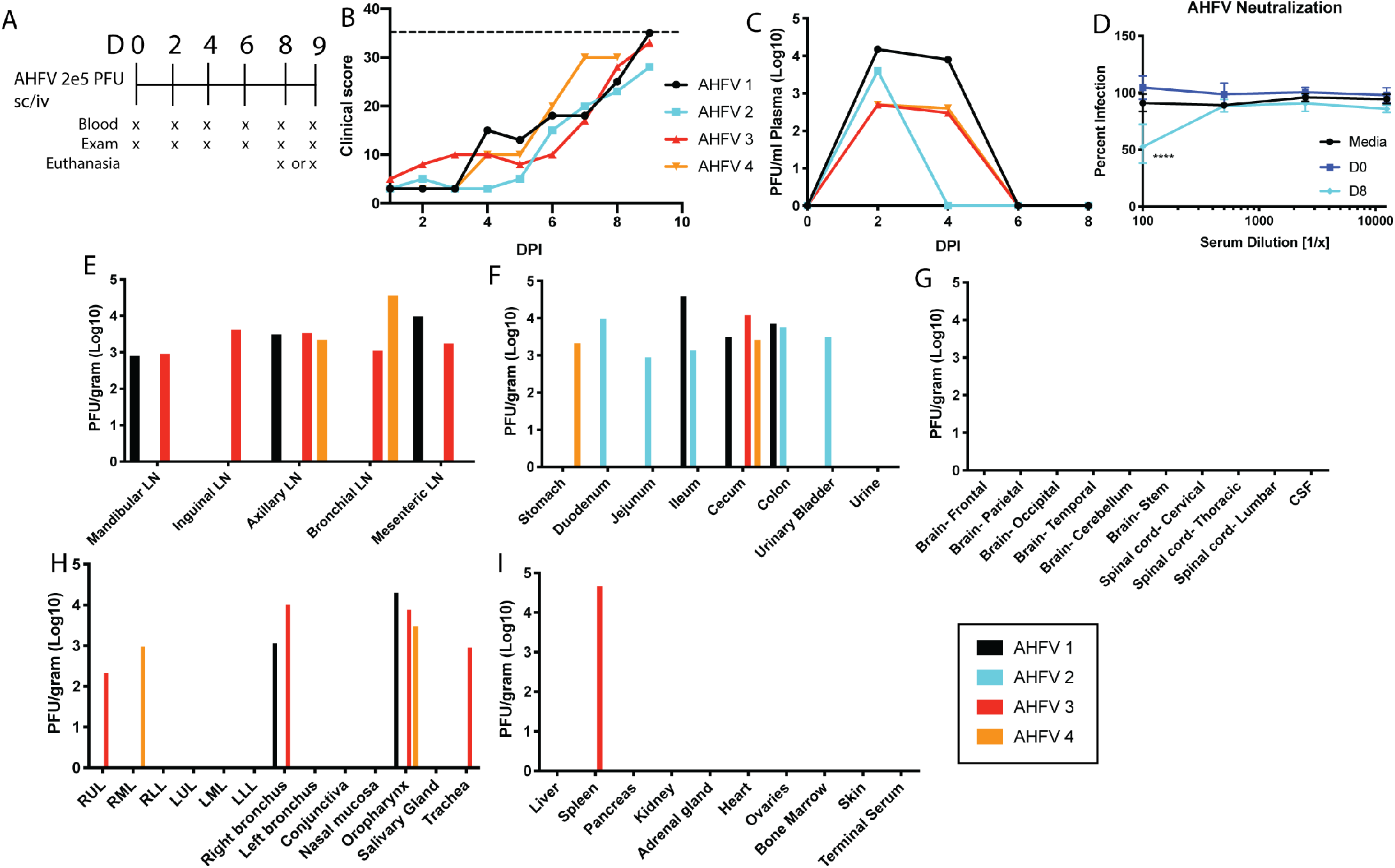
Pigtailed macaques are susceptible to disease caused by AHFV. Four pigtailed macaques (AHFV 1-4) were infected with 2 × 10^5^ pfu AHFV 200300001 sc/iv and monitored for clinical illness. **A**. Schedule for blood draws and exams in AHFV-infected animals. **B**. Clinical scores in four pigtailed macaques following infection with AHFV. Dotted line represents a score of 35. **C**. Viremia was measured in plasma over the first 8 days of infection by plaque assay. **D**. Neutralization assays were performed by mixing dilutions of heat-inactivated serum collected at the indicated times post infection with 100 pfu AHFV. Plaques were counted and normalized to D0. Media indicates no sera. Error bars show the SD across four animals. Statistics were performed using Sidak’s multiple comparison test, comparing D0 to D8 at all serum dilutions (*N* = 4; \*\*\*\**P* < 0.0001). **E-I**. At day 8 or 9 post infection, virus present in the tissue homogenate was measured by limiting dilution plaque assay, and values were normalized to 1 gram of tissue. Data from each individual animal are plotted.

### Changes in serum inflammatory markers and clinical blood parameters following infection with KFDV and AHFV

To measure the changes in inflammatory biomarkers following infection with KFDV or AHFV, serum cytokine and chemokines were measured throughout infection. Significant increases were observed with IL-6, MCP-1, IL-1RA, I-TAC, and IFNγ in the KFDV sc/iv group (Figure 3A-C, E), with peaks at 2, 4, or 6 days post infection. The same parameters were elevated in AHFV-infected animals, but only MCP-1 and IL-1RA reached statistical significance. Both KFDV sc/iv and AHFV infected animals had a significant decrease in IL-12 (Figure 3D). Other cytokines such as IFNγ, FGF-Basic, HGF, MIG, and Eotaxin were not significantly altered for all animals within the groups, but specific animals experienced changes in those cytokines following infection (Figure 3F-J). Mild changes in the reported cytokines and chemokines were also observed in the KFDV sc group. These animals had reduced IL-12 compared to baseline, but no other consistent changes were observed, likely due to the smaller group size and lower virus replication dynamics compared to the sc/iv groups. Statistics were not performed after day 6 due to the decreasing group sizes in the KFDV and AHFV groups.

**Figure 3.**
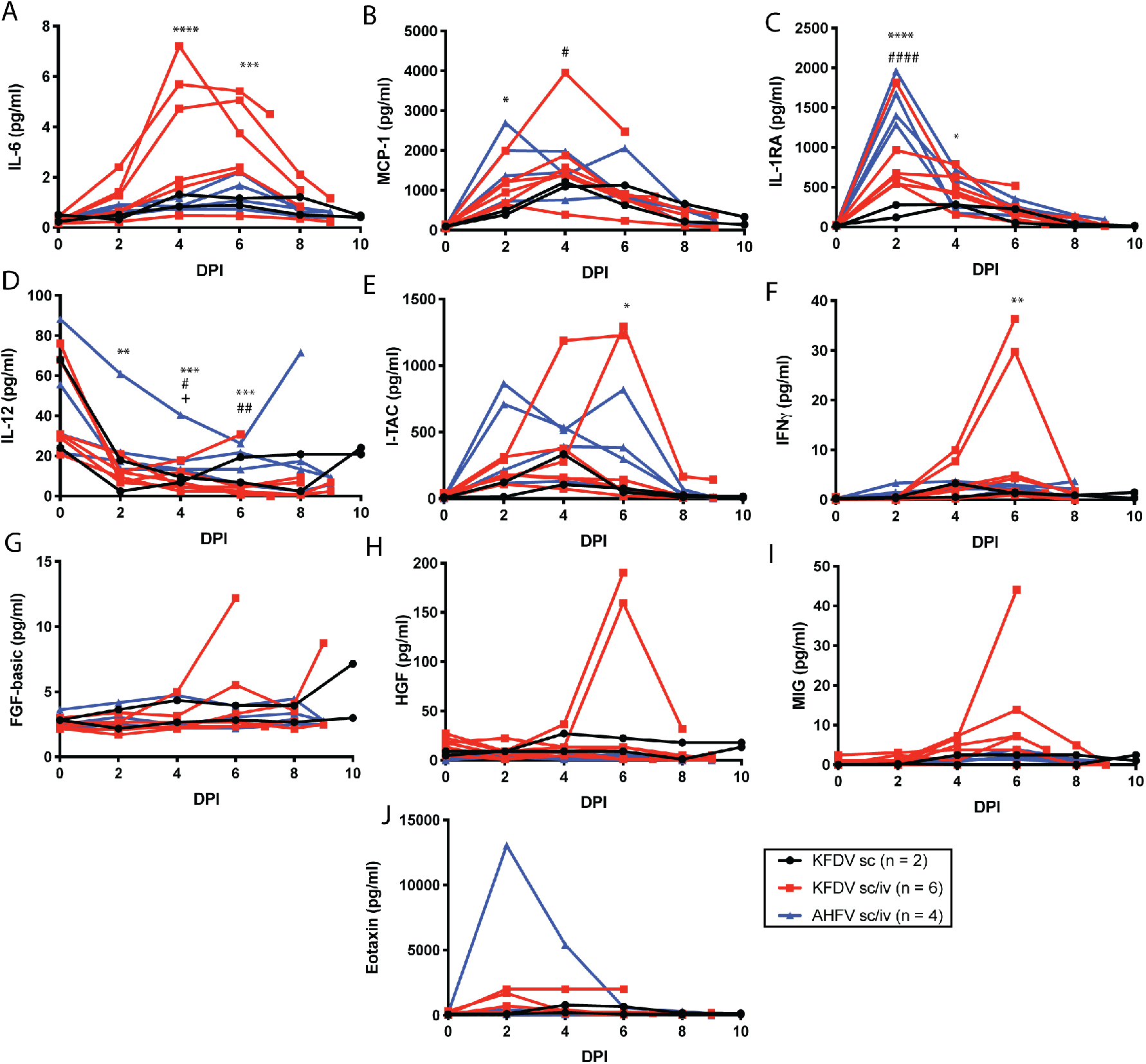
Cytokine and chemokine analysis in serum of KFDV and AHFV-infected animals. Pigtailed macaques were infected with KFDV by the sc or sc/iv route, or AHFV by the sc/iv route. The number of animals in each group is indicated in the legend. Serum was isolated from blood samples obtained during clinical exams at the indicated days post infection. Cytokine and chemokines were measured from sera using a 29-plex Monkey cytokine detection kit. The following parameters are included: **A**. IL-6, **B**. MCP-1, **C**. IL-1RA, **D**. IL-12, **E**. I-TAC, **F**. IFNγ, **G**. FGF-Basic, **H**. HGF, **I**. MIG, **J**. Eotaxin. Statistics were performed by comparing the change in the parameter compared to the D0 measurement using Sidaks multiple comparison test. Significance symbols are represented as follows: KFDV sc/iv *, AHFV sc/iv #, KFDV sc +. (KFDV sc/iv *N* = 6, AHFV sc/iv *N* = 4, KFDV sc *N* = 2; \**P* < 0.05, \*\**P* < 0.005, \*\*\**P* < 0.0005, \*\*\*\**P* < 0.0001.

Within the hemogram, all groups showed a decrease in HCT, HGB, and RBC throughout the study (Figure 4A-C). In some animals, these decreases resulted in a non-regenerative anemia particularly within the KFDV sc/iv group. Similarly, within the leukogram, all groups had a decrease in leukocytes that was driven by both significant decreases in neutrophils and lymphocytes (Figure 4D-F). The majority of the animals had leukopenia between D2 – 8 with some signs of recovery prior to the endpoint. The majority of the animals had neutropenia and lymphopenia during that same period. These findings are consistent with a previous study within bonnet macaques (*Macaca radiata*) (19). Animals in all groups demonstrated decreases in platelets throughout the study as well with some animals in the KFDV sc/iv and AHFV sc/iv becoming thrombocytopenic (<100 K/ul) (Figure 4G). The combination of the leukopenia and thrombocytopenia is a hallmark of human infection with KFDV (26).

**Figure 4.**
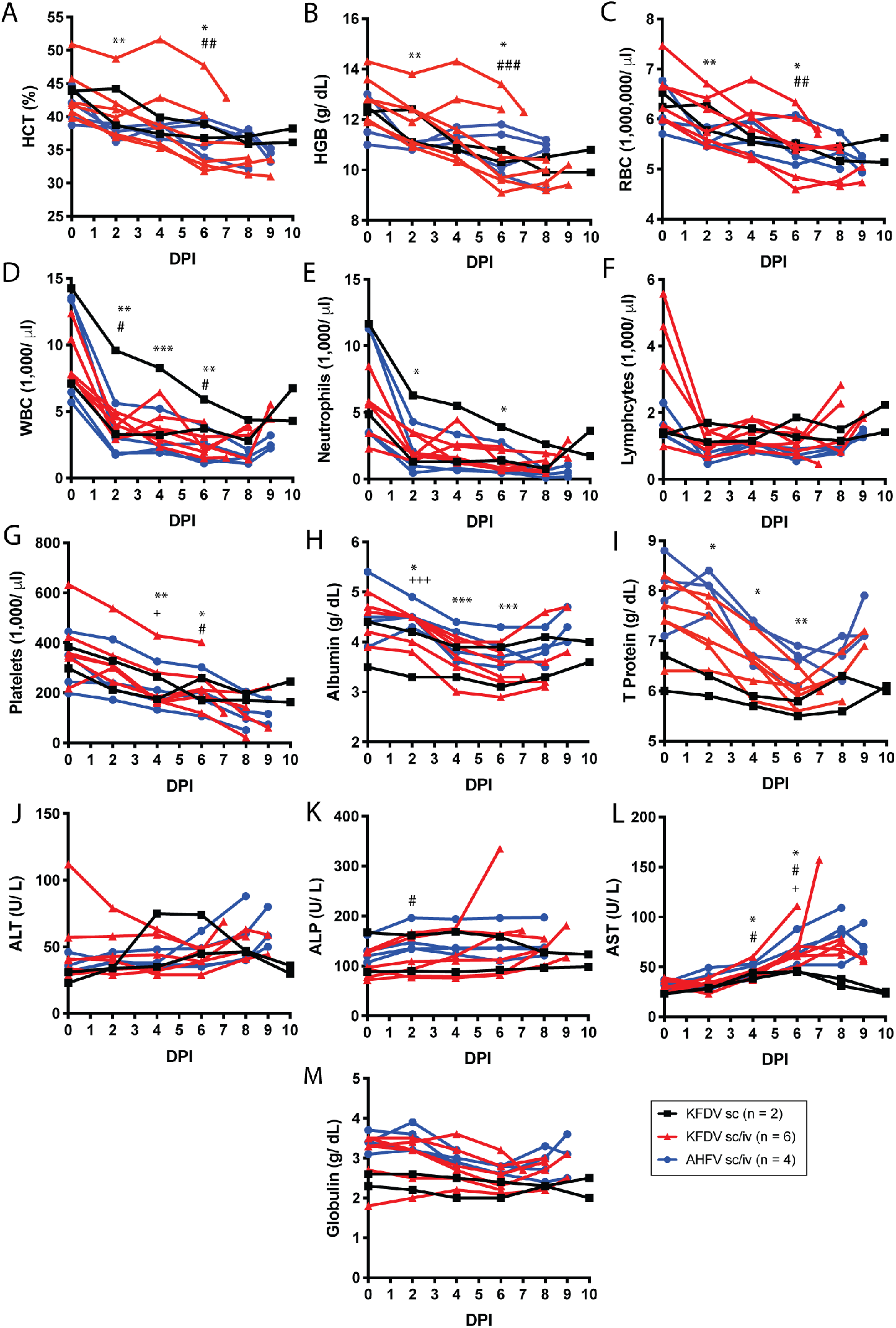
Blood parameters in pigtailed macaques infected with KFDV or AHFV. Blood was collected during exams as indicated. Complete blood counts were analyzed from EDTA blood (A-G), and blood chemistries were measured from sera (H-M). **A**. Hematocrit (HCT), **B**. Hemoglobin (HGB), **C**. Red blood cells (RBC), **D**. White blood cells (WBC), **E**. Neutrophils, **F**. Lymphocytes, **G**. Platelet counts, **H**. Albumin, **I**. Total protein, **J**. Alanine aminotransferase (ALT), **K**. Alanine phosphatase (ALP), **L**. Aspartate transaminase (AST), **M**. Globulin. Statistics were performed by comparing the change in the parameter at day 2, 4, and 6 compared to the day 0 measurement using Dunnett’s multiple comparison test. Significance symbols are represented as follows: KFDV sc/iv *, AHFV sc/iv #, KFDV sc +. (KFDV sc/iv *N* = 6, AHFV sc/iv *N* = 4, KFDV sc *N* = 2; \**P* < 0.05, \*\**P* < 0.005, \*\*\**P* < 0.0005, \*\*\*\**P* < 0.0001).

Although reported in humans and bonnet macaques that liver enyzmes increase with disease, this was not as clear within this model. In the AHFV sc/iv and KFDV sc/iv groups, a slight trend of increasing ALT and AST is noted particularly towards the later timepoints (Figure 4J-L). However these altered values are considered mild. More consistent were decreases in albumin, total protein and globulin up to D6 with signs of recovery after that timepoint (Figure 4H, I, M). Three KFDV sc/iv animals and one KFDV sc animal had hypoalbuminemia (< 3.5 g/dl) during the course of the study. The mouse model of KFDV has also been reported to show hypoalbuminemia in the later stage of disease (12).

### Histopathological findings in KFDV- and AHFV-infected PTMs

Gross pathology from KFDV- and AHFV-infected animals at necropsy is listed in Table 1. All six animals infected with KFDV sc/iv had pale or discolored livers. Two animals had hyperemic lungs with one animal having congested lungs and foam in the trachea. In the AHFV sc/iv group, two monkeys had enlarged spleens and/or livers, and two had dried blood in the nasal cavity. AHFV 1, the monkey reaching a clinical score of 35, had hemorrhages present on the liver and lungs. Thus, PTMs can be used to model both KFDV and AHFV disease, with clinical signs closely resembling human infection.

Despite some animals reaching the pre-established euthanasia criteria, it was surprising that no major histopathological findings could be definitively attributed to infection with KFDV or AHFV. In the KFDV sc/iv group, KFDV 4-8 had a chronic, marked-severe fatty infiltration of the liver parenchyma. KFDV 3-6 had minimal-mild amounts of lymphocytic infiltrates within the lamina propria of intestinal tract. In the AHFV sc/iv group, AHFV 1 and 4 had a chronic, marked-severe fatty infiltration of the liver parenchyma, and AHFV 3 had mild amyloidosis of the liver. AHFV 1 had a marked neutrophilic bronchopneumonia located in the left middle lung lobe and a minimal lesion in the left upper lobe. AHFV 4 had multifocal mucosal necrosis of the ileum. Within the KFDV sc group, KFDV 2 had moderate amyloidosis of the liver and minimal to mild lymphocytic infiltrates within the lamina propria of the intestinal tract. However, it is possible that these conditions are associated with chronic illness due to age and history of these individuals, and not directly due to virus infection.

### In-situ hybridization and Immunohistochemistry

In situ hybridization (ISH) was performed using a 20-nucleotide probe targeting KFDV NS4B and NS5. The target region of this probe in the KFDV P9605 genome is 93% identical to AHFV (200300001), and was capable of detecting AHFV genome. Within the KFDV sc inoculated group, only the bronchial lymph node of KFDV 1 showed positivity. However, in the KFDV sc/iv and AHFV sc/iv groups, immunoreactivity was observed in gut-associated lymphoid tissue (GALT), splenic, mediastinal, and mesenteric lymph node follicles (Figure 5). Consistent with the virus isolation data, KFDV positivity was also observed in the bronchus-associated lymphoid tissue (BALT) (Figure S3A). IHC was performed in the brain of KFDV 3 since infectious virus was directly isolated from brain samples of only this animal (Figure 1G). KFDV 3 had a focus of positivity in a cluster of mononuclear cells in the choroid plexus (Figure S3B). These data indicate that viral RNA in both KFDV and AHFV-infected animals is frequently associated with lymphoid tissue at the most acute stages of disease.

**Figure 5.**
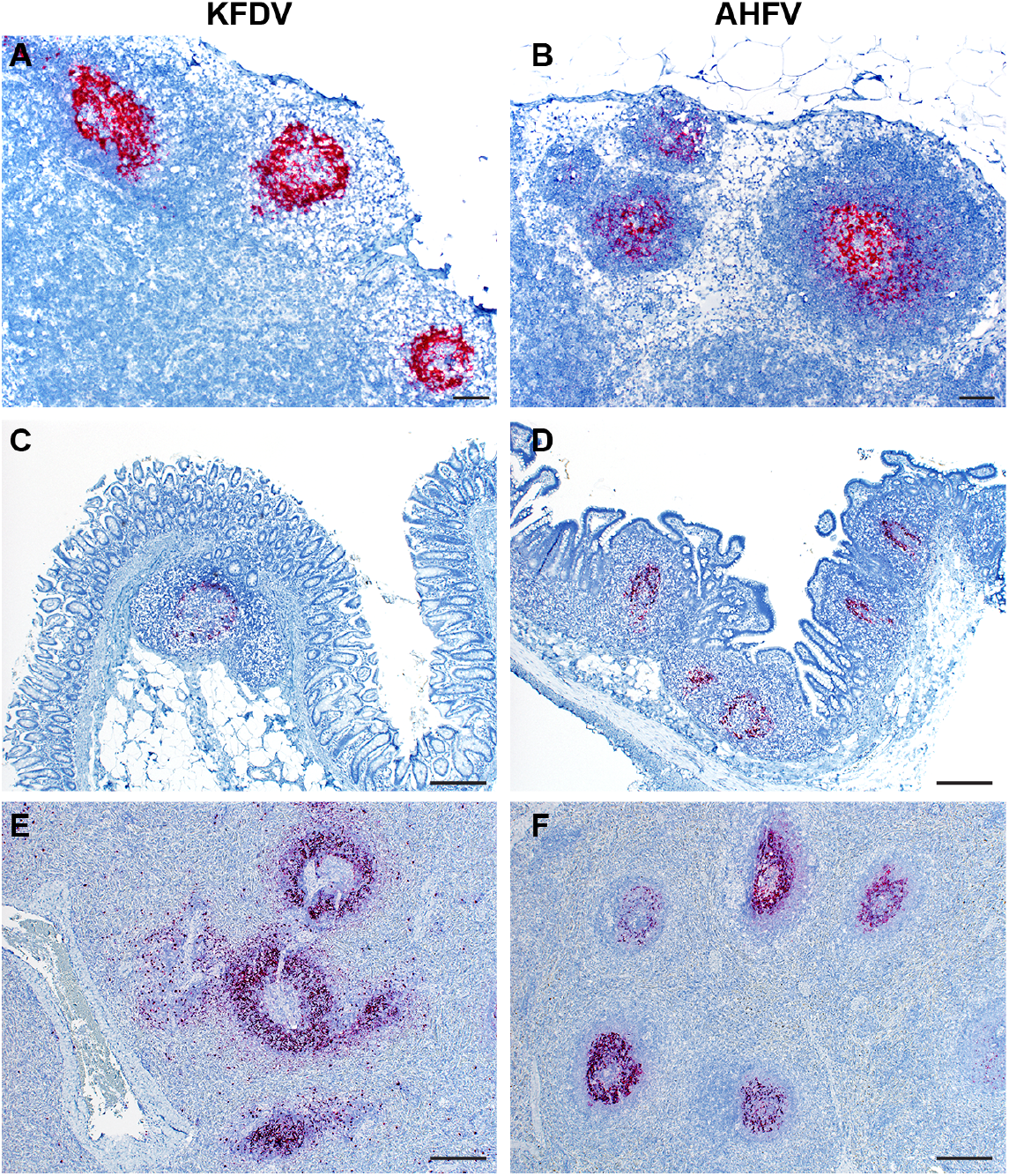
KFDV and AHFV RNA detection by ISH in the lymph node, intestine, and spleen of infected animals. A KFDV RNAscope probe was used to detect viral RNA from tissue sections derived from either KFDV or AHFV-infected PTMs. Tissues from KFDV sc/iv infected animals (**A, C, E**) or AHFV sc/iv infected animals (**B, D, F**) were examined. Representative images are displayed for **A, B**. mesenteric lymph node (40X, bar = 200 µm); **C**. cecum (100X, bar = 50 µm), **D**. ileum (100X, bar = 50 µm), and **E, F**. spleen (100X, bar = 50 µm).

Tissues were examined by ISH combined with IHC to directly identify cell types commonly associated with viral RNA positivity. Mediastinal lymph nodes were probed with antibodies directed towards CD3+, CD20+, or CD68+ cells for detection of T cells, B cells, and macrophages, respectively. KFDV RNA did not colocalize with CD3+ T cells. However, CD20+ B cells and CD68+ macrophages were associated with KFDV RNA (Figure 6). KFDV RNA association with CD20+ cells may either be direct infection of B cells, or the virus may be trapped on surface of B cells and/or follicular dendritic cells. Macrophages, a known flavivirus target, were also associated with KFDV genome positivity. Thus, the IHC and ISH provides evidence of infection in macrophages and strong association with B cell follicles.

**Figure 6.**
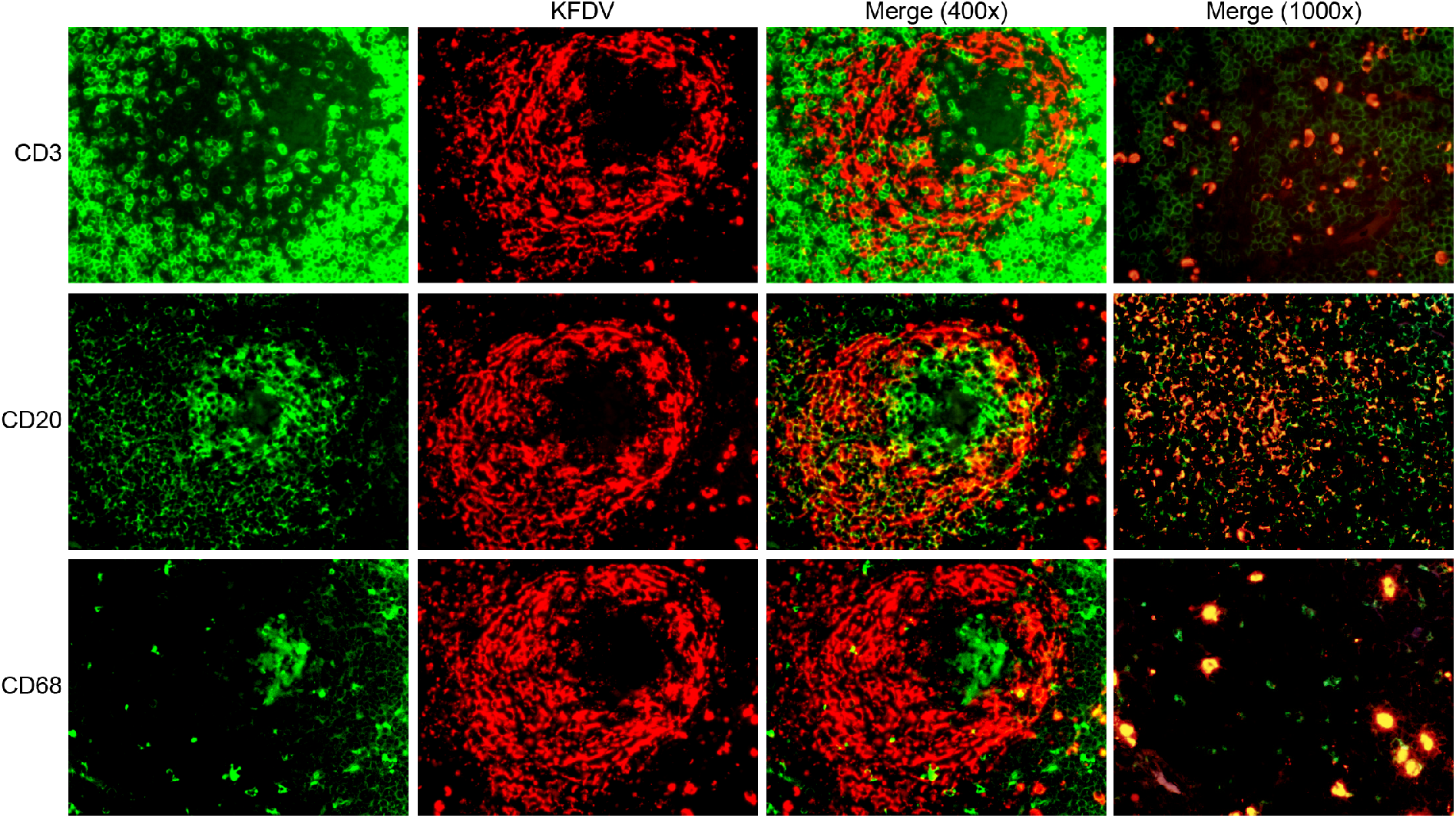
Mediastinal lymph node double staining with KFDV and CD3, CD20, or CD68. Mediastinal lymph node sections from an animal infected sc/iv with KFDV was subjected to RNAscope and IHC for CD3, CD20, and CD68 using the RNAscope VS Universal ISH-IHC HRP fluorescent assay. KFDV (red) and cell markers (green): CD3 (top row), CD20 (middle row), or CD68 (bottom row).

## Conclusion

The emergence of AHFV in the Arabian Peninsula and the increasing geographical range of KFDV in India highlights the urgent need for relevant animal models for testing vaccines and therapeutics. In this study, PTMs were tested for susceptibility and disease following infection with KFDV or AHFV. We found that animals infected by the sc/iv route demonstrated moderate to severe disease with KFDV and AHFV. KFDV infection in humans is commonly characterized by a sudden onset of fever, muscle weakness, back ache, vomiting, diarrhea, headache, reddened eyes, and some hemorrhagic manifestations such as bleeding from gums, or recurring epistaxis (29). In rare cases, the bleeding can be more severe, with hematemesis and hematochezia, and/or hemorrhage in the lungs. Consistent with case reports in humans, PTMs infected with KFDV had clinical signs such as weakness, loss of appetite, flushed appearance, epistaxis, and dehydration. Clinical signs were similar after infection with AHFV. One major advantage of the PTM model over current models is that these animals experienced marked thrombocytopenia and hemorrhagic manifestations following infection with KFDV and AHFV. In the KFDV-infected animals, most animals experienced epistaxis during the acute phase, except one animal that reached the euthanasia criteria at an early time point. Two KFDV-infected animals had hyperemic lungs, and one AHFV-infected animal had hemorrhages in the lung and liver. Together the PTM model for KFDV and AHFV closely resembles human infection and can be used for exploration into mechanisms of TBFV pathogenesis.

Infectious virus was frequently isolated from the gastrointestinal tissues, lungs, and lymph nodes in KFDV or AHFV-infected PTMs. In-situ hybridization revealed that the infection at these sites was associated with lymphoid tissue, particularly in the GALT, BALT, and the B cell zone within lymph nodes. These results are similar to reports of KFDV-infected bonnet macaques where antigen was seen in lymphocytes in the spleen as well as lymphoid cells in the gastrointestinal tract in KFDV-infected animals (18). However, KFDV-infected bonnet macaques showed virus positivity in epithelial cells within the gastrointestinal tract, but this was not observed in tissues from infected PTMs. Virus infection within lymphoid tissue such as the GALT or the BALT in NHPs has been shown with other viruses known to infect lymphocytes such as measles virus, and HIV (30, 31). Within lymphoid tissue, virus positivity in B cell follicles has been observed in other viral infection models including Zika virus-infected rhesus macaques, and in SIV-infected rhesus macaques (32, 33). This phenomenon could be evidence of trapped virus within B cell zones, possibly associated with follicular dendritic cells. While monocytes/macrophages are known targets of flavivirus infections, the consequences of potential B cell infection or antigen positivity is not described. A recent report of human infection with KFDV showed decreased levels of activated B cells in the acute phase of illness (34). The association of virus and B cells in the PTMs could provide mechanism for this phenomenon. B cell activation was not measured in the current study, but the KFDV-infected PTMs developed neutralizing antibodies associated with the resolution of viremia at 7-9 dpi, suggesting that the B cells were functional in this model.

KFDV is considered primarily a viscerotropic disease, where the second disease phase with CNS involvement is rare in humans (3, 29). Approximately three weeks following the acute phase, a second phase of illness may occur in roughly 10-20% of cases, characterized by recurrence of fever, headache, tremors, and neck stiffness (29, 35, 36). This study examined the KFDV and AHFV infection in PTMs up to 9 dpi in the animals inoculated sc/iv. Future work could extend the observation period to 4-5 weeks post infection to determine the incidence of this secondary phase of disease in PTMs. In KFDV-infected PTMs, virus was found in the brain of only one animal euthanized at 6 dpi, suggesting that virus localization in the brain is an uncommon event during the acute phase of disease. In contrast with the PTM model, KFDV is commonly isolated from the brain of infected bonnet macaques, even at acute time points (18-20). It should be noted that the animals used in this study were all females, 10-16 years old. These animals had a number of comorbidities which may have compounded the severity of illness, and additional studies will be necessary to determine if sex or age could influence the infection outcome.

Previous studies from the 1950s and 1960s attempting to develop models of KFDV in rhesus macaques described a development of viremia and neutralizing antibodies without overt disease, and the clinical descriptions of this model are absent or minimal (3, 16). Our current work suggests that PTMs may be more susceptible to infection with KFDV compared to historical reports in rhesus macaques. PTMs were successfully utilized as a model of HIV-1 in part because the genome of PTMs does not contain a restrictive allele of tripartite motif protein 5 (TRIM5) (27). TRIM5 from old-world monkeys including rhesus macaques imparts a strong block to replication of HIV by binding to the viral capsid lattice (37, 38). However, the PTM TRIM5 gene has undergone a retrotransposition event where cyclophilin A (CypA) replaces the domain of TRIM5 primarily responsible for capsid interactions (the B30.3 or PRY/SPRY domain) (39). We recently demonstrated that rhesus TRIM5 is also strongly restrictive to TBFVs, including KFDV and TBEV, through its interaction with the viral helicase NS3 (40). This restriction of TBFVs requires a functional SPRY domain, where TRIM-CypA fusions are not sufficient for restriction of TBFVs. Our ongoing hypothesis is that the pigtailed macaques may be more susceptible to infection with KFDV and AHFV in part because they possess a non-restrictive TRIM-CypA fusion protein. To develop the HIV-1 PTM model, it was necessary to overcome additional innate immune barriers such as APOBEC3, tetherin, and MX2 (27, 41, 42). Future investigation will determine what additional innate immune barriers to infection may exist for TBFVs, as was found with HIV-1.

One aspect of TBFV biology that was not addressed in this model was the ability of tick saliva to enhance TBFV dissemination and disease. Tick saliva contains a mixture of many factors known to modulate innate and adaptive immune responses (reviewed in (43, 44)). Rodents inoculated with virus mixed with tick salivary gland extract were more susceptible to paralysis after infection with a related flavivirus, Powassan virus (45). This and other studies demonstrate that tick saliva enhances pathogen transmission, dissemination, and impacts disease in sensitive animals. Tick saliva has not been used in TBFV NHP models, but its use may improve model development as it is clear from our pilot experiment with the sc route that the skin represents a strong barrier to systemic infection that was overcome by iv inoculation in order to develop PTMs as a disease model.

While no antiviral drugs are available to treat TBFV infections, several vaccines have been developed against TBFVs. The current vaccine for KFDV is a formalin-inactivated virus preparation that requires multiple boosters and has limited efficacy. Unfortunately, patients who received the KFDV vaccine may still develop viremia and clinical illness after KFDV infection (34, 46, 47). No vaccine has been approved for prevention of disease caused by AHFV. Thus, the PTM model of KFDV and AHFV pathogenesis will be useful for future efforts in vaccine and antiviral development, as well as determining mechanisms of virus pathogenesis.

## Methods

### Ethics statement

All work with infectious TBEV, KFDV, and AHFV was conducted at Biosafety Level 4 facilities at the Integrated Research Facility at the Rocky Mountain Laboratories (Hamilton, MT). All experiments involving PTMs were performed in strict accordance with the approved Institutional Animal Care and Use Committee protocol 2018-066-E and following recommendations from the Guide for the Care and Use of Laboratory Animals of the Office of Animal Welfare, National Institutes of Health and the Animal Welfare Act of the US Department of Agriculture, in an Association for Assessment and Accreditation of Laboratory Animal Care International (AAALAC)-accredited facility.

### Pigtailed Macaques

Twelve female PTMs aged 10 to 16 years of age were housed in a humidity, temperature, and light controlled facility in adjacent cages for social interaction. Animals were closely monitored at least twice daily by trained personnel. Animals were fed a commercial monkey chow twice per day, and their diets were supplemented with a variety of fruits, vegetables, and treats. Water was available ad libitum. Animals were provided with manipulanda, human interaction, and audio and visual enrichment. Virus inoculations were performed with the indicated virus diluted in sterile PBS. For subcutaneous (sc) infections, 1×10^5^ pfu was delivered in a single injection between the shoulder blades (1 ml volume). Intravenous (iv) inoculations were performed with 1×10^5^ pfu diluted virus using an IV catheter (1 ml volume). Animals were closely monitored and scored twice daily after showing signs of illness. Exams occurred on 0, 2, 4, 6, 8, 10, 14, 21, 28, 35, and 42 or 43 dpi. At each exam, blood was drawn in EDTA and serum separation tubes, and rectal and oral swabs were collected. Hematology analysis was completed on a ProCyte DX (IDEXX Laboratories, Westbrook, ME, USA) and the following parameters were evaluated: red blood cells (RBC), hemoglobin (Hb), hematocrit (HCT), mean corpuscular volume (MCV), mean corpuscular hemoglobin (MCH), mean corpuscular hemoglobin concentration (MCHC), red cell distribution weight (RDW), platelets, mean platelet volume (MPV), white blood cells (WBC), neutrophil count (abs and %), lymphocyte count (abs and %), monocyte count (abs and %), eosinophil count (abs and %), and basophil count (abs and %). Serum chemistries were completed on a VetScan VS2 Chemistry Analyzer (Abaxis, Union City, CA) and the following parameters were evaluated: glucose, blood urea nitrogen (BUN), creatinine, calcium, albumin, total protein, alanine aminotransferase (ALT), aspartate aminotransferase (AST), alkaline phosphatase (ALP), total bilirubin, globulin, sodium, potassium, chloride, and total carbon dioxide.

### Cells and viruses

Vero E6 cells (ATCC CCL-81) were grown in Dulbecco’s modified Eagle media (DMEM) containing 10% fetal bovine serum (FBS) and 1% antibiotics Penicillin-Streptomycin in an incubator at 37 °C and 5% CO_2_. KFDV (P9605), AHFV (200300001), and TBEV (Sofjin) seed stocks were obtained from the University of Texas Medical Branch (UTMB). Virus stocks were propagated on Vero cells, and passage 1 virus stocks (KFDV and AHFV) and passage 2 or 3 virus stocks (TBEV) were aliquoted and frozen in liquid nitrogen tanks.

### Virus isolation

Whole blood was collected in Vacutainer EDTA Tubes and overlayed on lymphocyte separation medium (Corning). After centrifugation, the plasma layer was collected and titrated on Vero cells by limiting dilution plaque assays. For virus isolation from tissues, samples were weighed and homogenized in DMEM containing 10% FBS and 1% antibiotics using 5 mm stainless steel beads (Qiagen) and the TissueLyser II (Qiagen). For plaque assays, samples were serially diluted in DMEM and added to Vero cells for 30 minutes to 1 hr while rocking the plate every 15 minutes. The cells were overlaid with 1.5% carboxymethylcellulose (Sigma) overlay in Minimum Essential Media (MEM). Following plaque development, plates were flooded with 10% formalin and plaques were visualized with a 1% crystal violet solution (4 days for KFDV and AHFV, 5 days for TBEV).

### Cytokine and chemokine analysis

Sera was inactivated by irradiation with 10 megarads (1 × 10^5^ Gray) according to standard biosafety protocols (48). Cytokines and chemokines were measured from irradiated sera using a magnetic Cytokine 29-Plex Monkey Panel (ThermoFisher) according to the manufacturer’s instructions. Sera was incubated with antibody beads for 2 hours followed by addition of detection and indicator antibodies. Cytokine levels were detected using a Bio-Plex 200 System with high-throughput fluidics (Bio-Rad).

### Neutralization assays

Neutralization tests were performed on Vero cells by a plaque reduction assay. Irradiated sera (10 megarads) were heat-inactivated for 1 hr at 56 °C. Five-fold serial dilutions of sera (1/50 to 1/6250) were prepared and mixed 1:1 with media containing 50 to 100 pfu of KFDV, TBEV, or AHFV for a final serum dilution of 1/100 to 1/12500. Virus and antibody complexes were incubated at 37 °C for two hours. The complexes were then plated onto Vero cells for 30 minutes followed by addition of CMC overlay. Following plaque development, Vero plates were flooded with 10% formalin and plaques visualized with crystal violet. The plaques were counted, and duplicate measurements with the same sera were averaged. The percent infection was then expressed as a percentage relative to the D0 serum.

### TBEV and KFDV RNA detection

RNA was extracted from 140 μl plasma using the QIAamp Viral RNA Mini Kit (Qiagen). RNA was irradiated (10 megarad) for removal from the BSL4 according to established sample removal protocols (48). Following removal, cDNA was generated using SuperScript VILO cDNA Synthesis Kit (ThermoFisher). Viral transcripts were detected by qRT-PCR using the Platinum Quantitative PCR SuperMix-UDG with ROX (ThermoFisher). Primer and probe sets used for qRT-PCR assays are as follows: KFDV forward primer: TGGCCAGCAGAGGGTTTTTA, reverse primer: AACGGCCCTCATGATGATCT, probe: CAAAGCGCAGGAGC. TBEV forward primer: GGCAGTCCCATCCTCAACTC, reverse primer: TGCTGACGTAGGTTTCATTGGT, probe: CTGTATGGGAACGGACTG.

### H&E

Tissues were fixed in 10% Neutral Buffered Formalin x2 changes, for a minimum of 7 days according to IBC approved protocols. Tissues were placed in cassettes and processed with a Sakura VIP-6 Tissue Tek, on a 12-hour automated schedule, using a graded series of ethanol, xylene, and PureAffin. Embedded tissues are sectioned at 5 μm and dried overnight at 42 °C prior to staining.

### In Situ Hybridization (ISH)

Chromogenic detection of KFDV viral RNA was performed on formalin fixed tissue using the RNAscope VS Universal AP assay (Advanced Cell Diagnostics Inc.) on the Ventana Discovery ULTRA stainer using an RNAscope® 2.5 VS Probe - V-KFDV-PP consisting of 20 probe pairs targeting the positive sense RNA at nt 7597-8486 of KFDV (Advanced Cell Diagnostics Inc. cat# 591199). ISH was performed according to manufacturer’s instructions. Fluorescent detection of KFDV ISH and IHC was performed using the RNAscope VS Universal ISH-IHC HRP fluorescent assay (Advanced Cell Diagnostics Inc.) on the Ventana Discovery ULTRA stainer according to manufacturer’s instructions. For the IHC, primary antibodies include CD3 (clone 2GV6) rabbit monoclonal (Roche Tissue Diagnostics cat#790-4341), CD20 rabbit polyclonal (Thermo Fisher Scientific cat#RB-9013), and CD68 (clone KP1) mouse monoclonal (Agilent cat#M081401-2). Secondary antibodies include Discovery Red 610 (Roche Tissue Diagnostics cat#760-245), Discovery FITC (Roche Tissue Diagnostics cat#760-232), Discovery OmniMap anti-Rabbit HRP (Roche Tissue Diagnostics cat# 760-4311), and Discovery OmniMap anti-mouse HRP (Roche Tissue Diagnostics cat# 760-4310). Slides were mounted using ProLong Diamond Antifade mountant w/ DAPI (Invitrogen cat# P36971).

### Statistics

Statistics were performed with a two-way ANOVA followed by Sidaks multiple comparison test or Dunnett’s multiple comparison test. Multiplicity-adjusted *P* values are reported, and statistical significance are indicated by \**P* < 0.05, \*\**P* < 0.005, \*\*\**P* < 0.0005, \*\*\*\**P* < 0.0001. Error bars represent the standard deviation. All statistical analysis were performed using Graphpad Prism.

## Acknowledgements

This research was supported by the Intramural Research Program of the NIH. We would like to acknowledge the Rocky Mountain Veterinary Branch animal caretakers for their assistance in these studies. We also acknowledge Heinz Feldmann for sharing of resources and his critical review of the manuscript.

---

**Figure S1.**
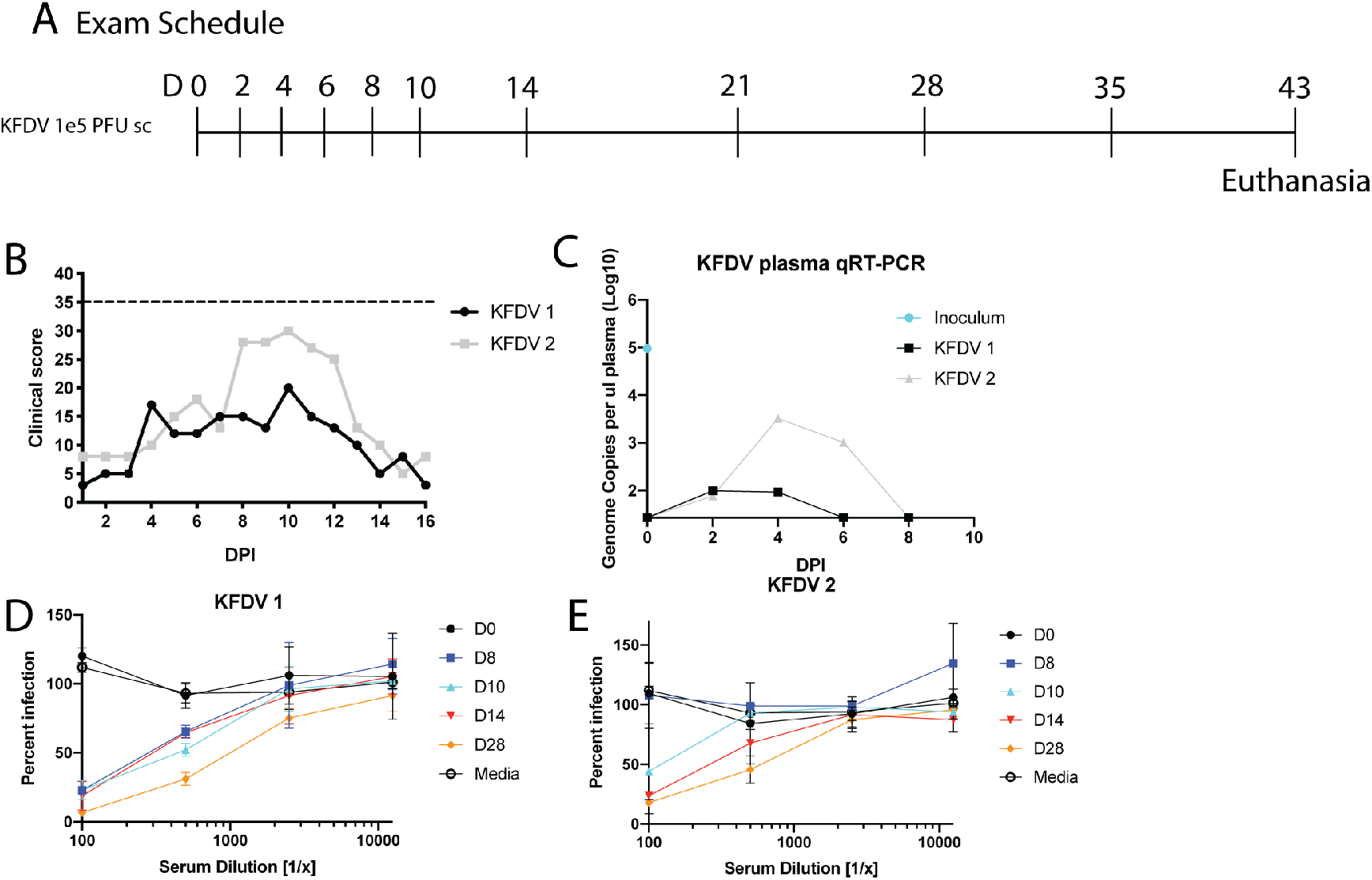
Pigtailed macaques infected with KFDV by the sc route. **A**. Two pigtailed macaques were infected with 10^5^ pfu KFDV (sc). Clinical exams occurred on D0, 2, 4, 6, 8, 10, 14, 21, 28, 35, and 43 post infection. Animals were euthanized at day 43 post infection. **B**. Clinical scores for KFDV-infected animals. **C**. Quantitative real-time PCR detection of KFDV transcripts derived from plasma on D0 through D8 post infection. **D-E**. Dilutions of heat-inactivated sera were tested for neutralization of KFDV by plaque assay. Data were normalized to the D0 values, and error bars represent the SD across duplicate measurements.

**Figure S2.**
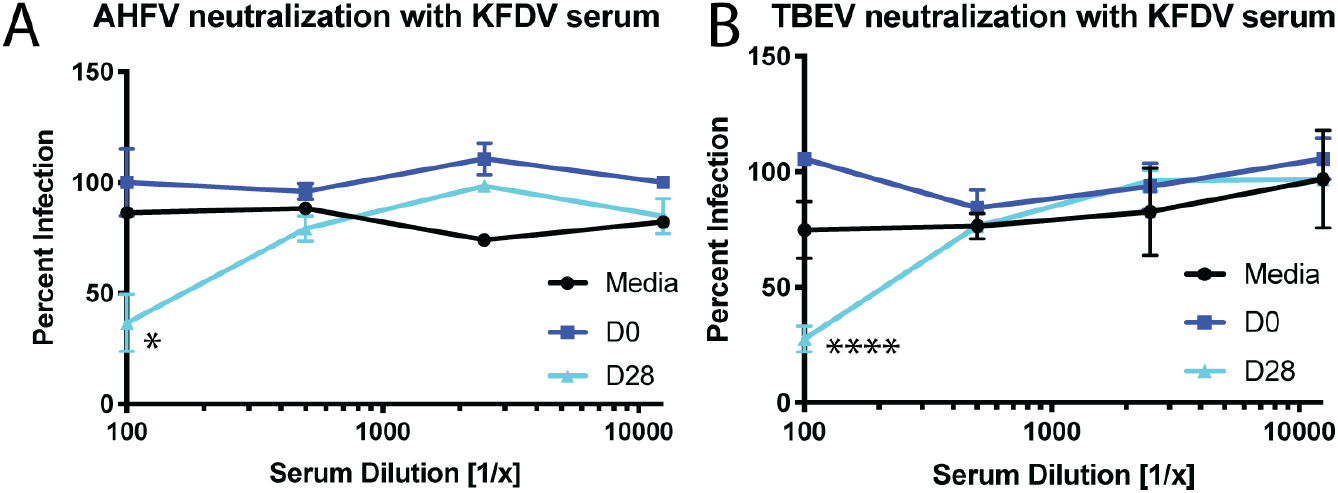
Sera from KFDV -infected animals cross-neutralizes AHFV and TBEV. Dilutions of heat-inactivated sera from KFDV 1 and KFDV 2 collected at 0 or 28 dpi were tested for neutralization of **A**. AHFV, and **B**. TBEV. Plaques were counted and normalized to the control D0 samples. Error bars represent the standard deviation across sera from two animals. Statistics were performed by comparing the change in the parameter compared to the D0 measurement using Sidaks multiple comparison test. (*N* = 2, \**P* < 0.05, **** *P* < 0.0001).

**Figure S3.**
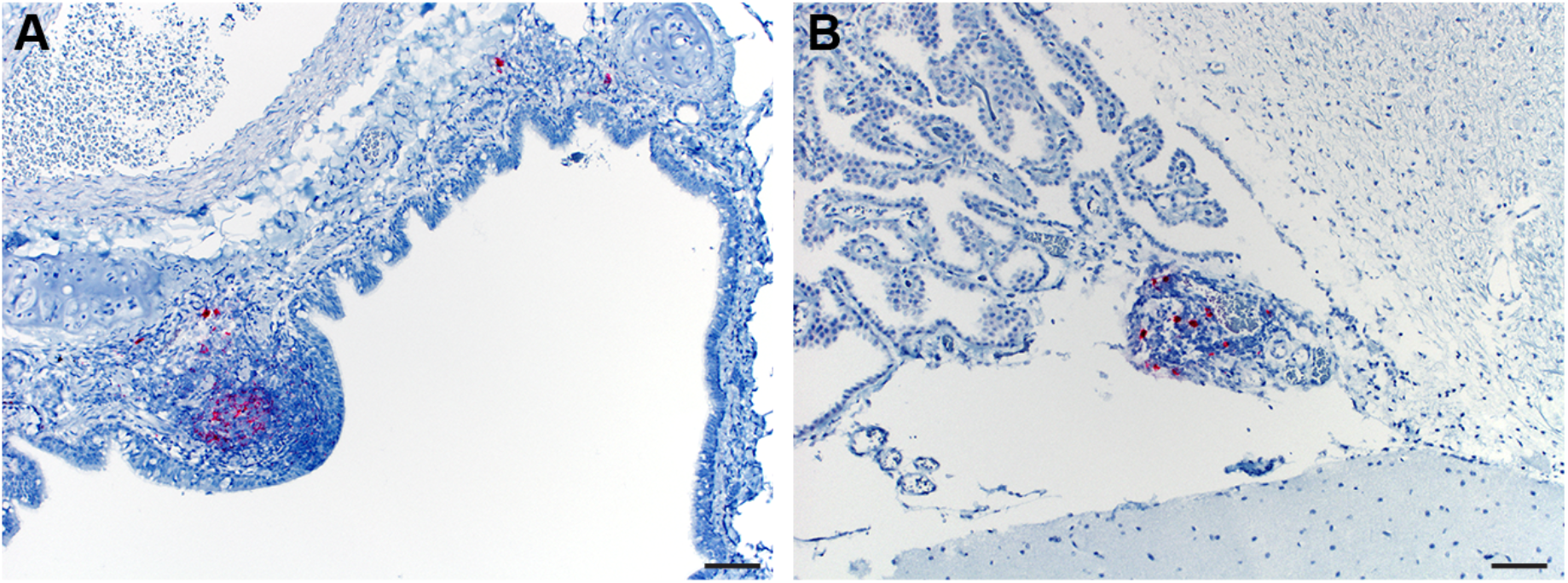
KFDV positivity in the lung and brain. Tissues from KFDV sc/iv infected animals were examined for immunoreactivity with a KFDV RNAscope probe for viral RNA. Immunoreactivity was identified in **A**. bronchus-associated lymphoid tissue (100X, bar = 50 µm), and **B**. choroid plexus (100X, bar = 50 µm).

**Table S1.** Clinical observations of KFDV-infected pigtailed macaques by the subcutaneous route.

|  | KFDV 1<br>(sc) | KFDV 2<br>(sc) | Frequency<br>(sc) n = 2 |
| --- | --- | --- | --- |
| Decreased appetite | Severe | Severe | 100% |
| Loss of interest in treats | N | N | 0% |
| Piloerection | Y | N | 50% |
| Hunched posture | Y | N | 50% |
| Loose stool | Y | N | 50% |
| No feces | N | N | 0% |
| clear nasal discharge | Y | Y | 100% |
| Transient epistaxis | N | Y | 50% |
| Unsteady | N | N | 0% |
| Slow and careful movements | N | N | 0% |
| Reluctant to move | N | N | 0% |
| Using cage for support | N | N | 0% |
| Dehydration | N | N | 0% |
| CRT 2 seconds or more | D10, 3 sec. | N | 50% |
| Temperature change > 1 °C | N | N | 0% |
| Flushed Appearance | N | N | 0% |
| Mild Facial edema | N | N | 0% |
| Other Comments |  |  |  |
| Score 35 | N | N | 0% |

## References

1. Chakraborty S, Andrade FCD, Ghosh S, Uelmen J, Ruiz MO. Historical Expansion of Kyasanur Forest Disease in India From 1957 to 2017: A Retrospective Analysis. Geohealth. 2019;3(2):44–55.

2. Munivenkatappa A, Sahay RR, Yadav PD, Viswanathan R, Mourya DT. Clinical & epidemiological significance of Kyasanur forest disease. Indian J Med Res. 2018;148(2):145–50.

3. Work TH. Russian spring-summer virus in India: Kyasanur Forest disease. Prog Med Virol. 1958;1:248–79.

4. Zaki AM. Isolation of a flavivirus related to the tick-borne encephalitis complex from human cases in Saudi Arabia. Trans R Soc Trop Med Hyg. 1997;91(2):179–81.

5. Dodd KA, Bird BH, Khristova ML, Albariño CG, Carroll SA, Comer JA, et al. Ancient ancestry of KFDV and AHFV revealed by complete genome analyses of viruses isolated from ticks and mammalian hosts. PLoS Negl Trop Dis. 2011;5(10):e1352.

6. Carletti F, Castilletti C, Di Caro A, Capobianchi MR, Nisii C, Suter F, et al. Alkhurma hemorrhagic fever in travelers returning from Egypt, 2010. Emerg Infect Dis. 2010;16(12):1979–82.

7. Madani TA, Azhar EI, Abuelzein e-T, Kao M, Al-Bar HM, Abu-Araki H, et al. Alkhumra (Alkhurma) virus outbreak in Najran, Saudi Arabia: epidemiological, clinical, and laboratory characteristics. J Infect. 2011;62(1):67–76.

8. Pugliese A, Beltramo T, Torre D. Emerging and re-emerging viral infections in Europe. Cell Biochem Funct. 2007;25(1):1–13.

9. Smura T, Tonteri E, Jääskeläinen A, von Troil G, Kuivanen S, Huitu O, et al. Recent establishment of tick-borne encephalitis foci with distinct viral lineages in the Helsinki area, Finland. Emerg Microbes Infect. 2019;8(1):675–83.

10. Tigabu B, Juelich T, Bertrand J, Holbrook MR. Clinical evaluation of highly pathogenic tick-borne flavivirus infection in the mouse model. J Med Virol. 2009;81(7):1261–9.

11. Sawatsky B, McAuley AJ, Holbrook MR, Bente DA. Comparative pathogenesis of Alkhumra hemorrhagic fever and Kyasanur forest disease viruses in a mouse model. PLoS Negl Trop Dis. 2014;8(6):e2934.

12. Dodd KA, Bird BH, Jones ME, Nichol ST, Spiropoulou CF. Kyasanur Forest disease virus infection in mice is associated with higher morbidity and mortality than infection with the closely related Alkhurma hemorrhagic fever virus. PLoS One. 2014;9(6):e100301.

13. Palus M, Vojtíšková J, Salát J, Kopecký J, Grubhoffer L, Lipoldová M, et al. Mice with different susceptibility to tick-borne encephalitis virus infection show selective neutralizing antibody response and inflammatory reaction in the central nervous system. J Neuroinflammation. 2013;10:77.

14. Basu A, Yadav P, Prasad S, Badole S, Patil D, Kohlapure RM, et al. An Early Passage Human Isolate of Kyasanur Forest Disease Virus Shows Acute Neuropathology in Experimentally Infected CD-1 Mice. Vector Borne Zoonotic Dis. 2016;16(7):496–8.

15. Nikiforuk AM, Tierny K, Cutts TA, Kobasa DK, Theriault SS, Cook BWM. Kyasanur Forest disease virus non-mouse animal models: a pilot study. BMC Res Notes. 2020;13(1):291.

16. Ilyenko VI, Pokrovskaya OA. Clinical picture in M. rhesus monkeys infected with various strains of tick-borne encephalitis virus. New York: Czechoslovak Academy of Sciences Praha; 1960. p. 266–9.

17. Shah KV, Dandawate CN, Bhatt PN. Kyasanur forest disease virus: viremia and challenge studies in monkeys with evidence of cross-protection by Langat virus infection. F1000Res. 2012;1:61.

18. Kenyon RH, Rippy MK, McKee KT, Zack PM, Peters CJ. Infection of Macaca radiata with viruses of the tick-borne encephalitis group. Microb Pathog. 1992;13(5):399–409.

19. Patil DR, Yadav PD, Shete A, Chaubal G, Mohandas S, Sahay RR, et al. Study of Kyasanur forest disease viremia, antibody kinetics, and virus infection in target organs of Macaca radiata. Sci Rep. 2020;10(1):12561.

20. Webb HE, Burston J. Clinical and pathological observations with special reference to the nervous system in Macaca radiata infected with Kyasanur Forest Disease virus. Trans R Soc Trop Med Hyg. 1966;60(3):325–31.

21. Webb HE, Chaterjea JB. Clinico-pathological observations on monkeys infected with Kyasanur Forest disease virus, with special reference to the haemopoietic system. Br J Haematol. 1962;8:401–13.

22. Adams Waldorf KM, Nelson BR, Stencel-Baerenwald JE, Studholme C, Kapur RP, Armistead B, et al. Congenital Zika virus infection as a silent pathology with loss of neurogenic output in the fetal brain. Nat Med. 2018;24(3):368–74.

23. Adams Waldorf KM, Stencel-Baerenwald JE, Kapur RP, Studholme C, Boldenow E, Vornhagen J, et al. Fetal brain lesions after subcutaneous inoculation of Zika virus in a pregnant nonhuman primate. Nat Med. 2016;22(11):1256–9.

24. Baskin CR, García-Sastre A, Tumpey TM, Bielefeldt-Ohmann H, Carter VS, Nistal-Villán E, et al. Integration of clinical data, pathology, and cDNA microarrays in influenza virus-infected pigtailed macaques (Macaca nemestrina). J Virol. 2004;78(19):10420–32.

25. Subekti DS, Tjaniadi P, Lesmana M, McArdle J, Iskandriati D, Budiarsa IN, et al. Experimental infection of Macaca nemestrina with a Toronto Norwalk-like virus of epidemic viral gastroenteritis. J Med Virol. 2002;66(3):400–6.

26. Lusso P, Crowley RW, Malnati MS, Di Serio C, Ponzoni M, Biancotto A, et al. Human herpesvirus 6A accelerates AIDS progression in macaques. Proc Natl Acad Sci U S A. 2007;104(12):5067–72.

27. Hatziioannou T, Ambrose Z, Chung NP, Piatak M, Yuan F, Trubey CM, et al. A macaque model of HIV-1 infection. Proc Natl Acad Sci U S A. 2009;106(11):4425–9.

28. VanBlargan LA, Errico JM, Kafai NM, Burgomaster KE, Jethva PN, Broeckel RM, et al. Broadly neutralizing monoclonal antibodies protect against multiple tick-borne flaviviruses. J Exp Med. 2021;218(5).

29. Work TH, Trapido H, Murthy DP, Rao RL, Bhatt PN, Kulkarni KG. Kyasanur forest disease. III. A preliminary report on the nature of the infection and clinical manifestations in human beings. Indian J Med Sci. 1957;11(8):619–45.

30. de Vries RD, McQuaid S, van Amerongen G, Yüksel S, Verburgh RJ, Osterhaus AD, et al. Measles immune suppression: lessons from the macaque model. PLoS Pathog. 2012;8(8):e1002885.

31. Thompson CG, Gay CL, Kashuba ADM. HIV Persistence in Gut-Associated Lymphoid Tissues: Pharmacological Challenges and Opportunities. AIDS Res Hum Retroviruses. 2017;33(6):513–23.

32. Brenchley JM, Vinton C, Tabb B, Hao XP, Connick E, Paiardini M, et al. Differential infection patterns of CD4+ T cells and lymphoid tissue viral burden distinguish progressive and nonprogressive lentiviral infections. Blood. 2012;120(20):4172–81.

33. Hirsch AJ, Smith JL, Haese NN, Broeckel RM, Parkins CJ, Kreklywich C, et al. Zika Virus infection of rhesus macaques leads to viral persistence in multiple tissues. PLoS Pathog. 2017;13(3):e1006219.

34. Devadiga S, McElroy AK, Prabhu SG, Arunkumar G. Dynamics of human B and T cell adaptive immune responses to Kyasanur Forest disease virus infection. Sci Rep. 2020;10(1):15306.

35. Webb HE, Rao RL. Kyasanur forest disease: a general clinical study in which some cases with neurological complications were observed. Trans R Soc Trop Med Hyg. 1961;55:284–98.

36. Adhikari Prabha MR, Prabhu MG, Raghuveer CV, Bai M, Mala MA. Clinical study of 100 cases of Kyasanur Forest disease with clinicopathological correlation. Indian J Med Sci. 1993;47(5):124–30.

37. Stremlau M, Owens CM, Perron MJ, Kiessling M, Autissier P, Sodroski J. The cytoplasmic body component TRIM5alpha restricts HIV-1 infection in Old World monkeys. Nature. 2004;427(6977):848–53.

38. Stremlau M, Perron M, Lee M, Li Y, Song B, Javanbakht H, et al. Specific recognition and accelerated uncoating of retroviral capsids by the TRIM5alpha restriction factor. Proc Natl Acad Sci U S A. 2006;103(14):5514–9.

39. Newman RM, Hall L, Kirmaier A, Pozzi LA, Pery E, Farzan M, et al. Evolution of a TRIM5-CypA splice isoform in old world monkeys. PLoS Pathog. 2008;4(2):e1000003.

40. Chiramel AI, Meyerson NR, McNally KL, Broeckel RM, Montoya VR, Méndez-Solís O, et al. TRIM5α Restricts Flavivirus Replication by Targeting the Viral Protease for Proteasomal Degradation. Cell Rep. 2019;27(11):3269-83.e6.

41. Hatziioannou T, Del Prete GQ, Keele BF, Estes JD, McNatt MW, Bitzegeio J, et al. HIV-1-induced AIDS in monkeys. Science. 2014;344(6190):1401–5.

42. Schmidt F, Keele BF, Del Prete GQ, Voronin D, Fennessey CM, Soll S, et al. Derivation of simian tropic HIV-1 infectious clone reveals virus adaptation to a new host. Proc Natl Acad Sci U S A. 2019;116(21):10504–9.

43. Kotál J, Langhansová H, Lieskovská J, Andersen JF, Francischetti IM, Chavakis T, et al. Modulation of host immunity by tick saliva. J Proteomics. 2015;128:58–68.

44. Wikel SK. Tick modulation of host immunity: an important factor in pathogen transmission. Int J Parasitol. 1999;29(6):851–9.

45. Hermance ME, Thangamani S. Tick Saliva Enhances Powassan Virus Transmission to the Host, Influencing Its Dissemination and the Course of Disease. J Virol. 2015;89(15):7852–60.

46. Kasabi GS, Murhekar MV, Sandhya VK, Raghunandan R, Kiran SK, Channabasappa GH, et al. Coverage and effectiveness of Kyasanur forest disease (KFD) vaccine in Karnataka, South India, 2005-10. PLoS Negl Trop Dis. 2013;7(1):e2025.

47. Kiran SK, Pasi A, Kumar S, Kasabi GS, Gujjarappa P, Shrivastava A, et al. Kyasanur Forest disease outbreak and vaccination strategy, Shimoga District, India, 2013-2014. Emerg Infect Dis. 2015;21(1):146–9.

48. Feldmann F, Shupert WL, Haddock E, Twardoski B, Feldmann H. Gamma Irradiation as an Effective Method for Inactivation of Emerging Viral Pathogens. Am J Trop Med Hyg. 2019;100(5):1275–7.

